# Australian *Culex annulirostris* mosquitoes are competent vectors for Japanese encephalitis virus genotype IV

**DOI:** 10.1101/2024.06.13.598393

**Authors:** Melissa J. Klein, Sarah A. Jackson, Willy W. Suen, Jean Payne, Darcy Beveridge, Mel Hargreaves, Donna Gillies, Jianning Wang, Kim R. Blasdell, Mike Dunn, Adam J. López-Denman, David T. Williams, Prasad N. Paradkar

## Abstract

Japanese encephalitis virus (JEV) is transmitted by *Culex* species of mosquitoes. In 2022, JEV belonging to a previously unrecognised lineage of genotype IV (GIV) caused a major outbreak of JE in South-eastern Australia, resulting in human cases and affecting piggeries. *Cx. annulirostris* has previously been implicated as the major vector of JEV in northern Australia where the virus has circulated since its first detection in 1995. Here, we showed that experimental infection of a laboratory colony of Australian *Cx. annulirostris* with the isolate JEV NSW/22 resulted in a 100% mosquito infection rate, with 87% of mosquito saliva samples testing positive by RT-qPCR at 14 days post-infection. Immunohistochemistry confirmed the presence of a replicating virus in the mosquito midgut and dissemination throughout the body, including the salivary glands. Our results also showed evidence of transovarial transmission of this virus; however, transstadial transmission from the eggs to the adult stage was not found. Comparison with an Indonesian isolate of GIV JEV and previous Australian isolates belonging to genotypes I and II showed that infection with JEV NSW/22 resulted in higher viral titres in the early stage of infection and higher proportions of mosquitoes with JEV-positive saliva, indicating a greater transmission potential compared to other isolates. This study provides compelling experimental evidence that Australian *Cx. annulirostris* is a highly efficient vector for the 2022 Australian JEV GIV outbreak strain.

## 1. Introduction

The mosquito-borne Japanese encephalitis virus (JEV; *Orthoflavivirus japonicum)* recently emerged in South-eastern Australia[1]. JEV is the major cause of human viral encephalitis in Southeast Asia, with an estimated 68,000 to 100,000 cases occurring annually[2,3]. JEV is also an important veterinary pathogen, where infections of horses can result in encephalitic disease with similar outcomes to human JE[4], while infection of pigs can lead to reproductive disease and production losses[5], although most infections remain asymptomatic. JEV is a zoonotic virus transmitted between species of *Culex* mosquitoes and vertebrate hosts. Ardeid water birds, including herons and egrets, are thought to be the principal wildlife reservoir and play a role in virus maintenance and long-range spread, while pigs are the major amplifying host[6,7]. Other vertebrate hosts such as bats and young poultry may also play a role in the virus transmission cycle, however their role is not fully understood [6,7].

Five genotypes (GI-GV) of JEV have been identified to date, with two separate lineages described for GI [8,9]. JEV is believed to have evolved from the Indo-Malaysia region, where all five genotypes have been detected, and subsequently dispersed to neighbouring regions where distinct patterns of distribution for each genotype have been found[8,10]. Co-circulation of one to three different genotypes is known to occur and genotype displacement has previously been documented [9,11]. In the Australasian region, JEV GII was first detected in the Torres Strait Islands of northern Australia following an outbreak of JE in 1995 [12] and in mosquitoes collected in the Western Province of Papua New Guinea (PNG) in 1997 and 1998 [13]. Genotype I was subsequently detected in the Torres Strait Islands in 2000 [13] and northern Queensland in Australia in 2004 [14] and later shown to belong to the G1a lineage [9].

In 2021, a fatal human case of JE occurred in a resident of the Tiwi Islands of the Northern Territory in Australia, caused by a novel lineage of GIV JEV[15,16]. This case proved to be the precursor of an unprecedented widespread outbreak of JE in Australia in 2022, which was detected for the first time in South-eastern Australia since surveillance began [1,17]. A total of 45 human cases of JE and more than 80 infected commercial piggeries were reported, with the majority of cases occurring in New South Wales, Victoria and South Australia [18]. JEV GIV is the third genotype to emerge in Australia and before its emergence in Australia, this genotype was rarely detected beyond a geographic range predominantly restricted to Indonesia [19,20]. The earliest mosquito isolates of GIV were derived from *Cx. tritaeniorhynchus, Cx. vishnui* or mixed species pools collected in 1980 and 1981 in Indonesia [20]. More recently, GIV JEV was isolated in 2017 from pigs in Bali, Indonesia [21] and in 2019 from *Cx. vishnui* mosquitoes[22] and a human travel case [23]. JEV GIV was also detected in Papua New Guinea *Cx. gelidus* in 2019 [1].

*Cx. tritaeniorhynchus* is considered the primary vector for JEV throughout Southeast Asia [24,25] with all 5 genotypes having been isolated from field samples of these mosquitoes and vector competence has been confirmed via laboratory experiments[26–28]. Previous studies have also determined several other species such as *Cx. vishnui* [29], *Cx. gelidus* [30], *Cx. pipiens* [31], *Cx. quinquefasciatus* [32], *Ae. albopictus* [33] *and Ae. japonicus* [27] to be competent vectors in laboratory settings. Furthermore, previous studies have also demonstrated vertical transmission of JEV in *Cx. tritaeniorhynchus* and *Ae. japonicus* [34,35]. In Australia, based on mosquito surveillance and host feeding patterns, *Cx. annulirostris* is considered the primary vector for JEV [36,37]. This species is widely distributed across Australia and commonly feeds on birds and pigs. Previous vector competence experiments using laboratory colonies have shown high transmission rates for JEV GII strain [32]. However, the vector competence of *Cx. annulirostris* for JEV GIV is not known. Given the prevalence of these mosquitoes in Australia, it is important to confirm the vector competence of Australian *Cx. annulirostris* to fully understand the risk this species presents for the continued circulation and spread of JEV GIV in Australia.

In this study, we conducted vector competence and vertical transmission experiments using a laboratory colony of Australian *Cx. annulirostris* and the Australian NSW/22 outbreak isolate. Experiments were also conducted using other historical Australian GI and GII isolates, as well as an early Indonesian isolate of GIV, to compare the infection dynamics and relative fitness of these viruses in *Cx. annulirostris*.

## 2. Materials and Methods

### 2.1 Mosquitoes

We performed experiments using a laboratory colony of *Cx. annulirostris* originating from Shepparton, Victoria. Mosquito egg rafts were hatched 4–5 days post blood meal and larvae were reared in small plastic trays with reverse osmosis water and fish food (Sera) ad libitum until the pupae stage. Emerging adults were housed in Plexiglas cages (Resi-Plex, Australia) with 10% sugar solution ad libitum which was removed 24 hours before a chicken blood meal using the Hemotek system (Hemotek). Blood was collected from specific pathogen-free chickens into lithium heparin vacutainers (Becton Dickinson) (ACDP Animal Ethics 1957, 22010). Mosquitoes were housed at 26.5+/-1°C, 65% relative humidity and 12-hour light:dark photoperiod and all experiments were performed under these conditions.

### 2.2 Viruses

Four isolates of JEV were used in the vector competence experiments: TS00 (GI), FU (GII), JKT6468 (GIV) and the recently emerged Australian isolate O-0883/NSW/2022 (NSW/22; GIV) [38]. JEV JKT6468 was isolated from *Cx. tritaeniorhynchus* mosquitoes collected in Flores, Indonesia in 1981 [19]. The JEV FU isolate was isolated from the blood of an infected resident of Badu Island during the first emergence of JEV in the Torres Straits in 1995[12], and GI isolate, TS00, was derived from blood collected from a pig on Badu Island in 2000 [39] (Table 1). Virus isolates were passaged in C6/36 cells (*Aedes albopictus*) and Vero cells (African green monkey kidney). The resulting viral supernatant for each isolate was clarified and titrated in Vero cells, with titres calculated using the Reed Muench method[40].

**Table 1:**
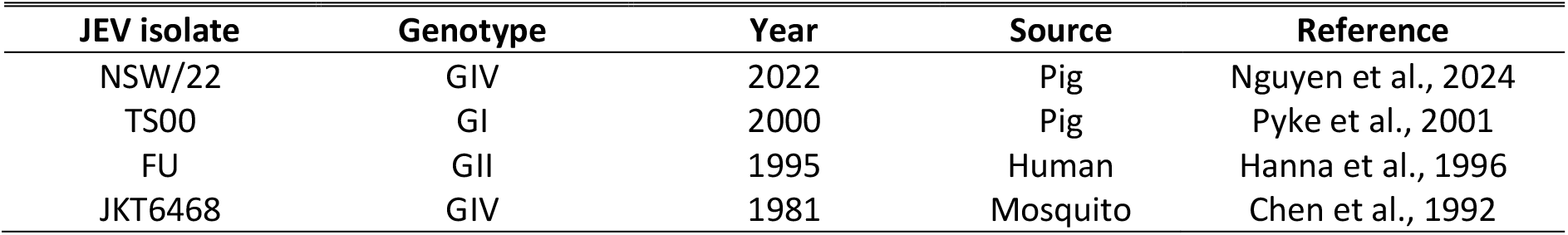
Japanese encephalitis virus isolates used in this study.

| JEV isolate | Genotype | Year | Source | Reference |
| --- | --- | --- | --- | --- |
| NSW/22 | GIV | 2022 | Pig | Nguyen et al., 2024 |
| TS00 | GI | 2000 | Pig | Pyke et al., 2001 |
| FU | GII | 1995 | Human | Hanna et al., 1996 |
| JKT6468 | GIV | 1981 | Mosquito | Chen et al., 1992 |

### 2.3 Vector competence

To assess vector competence, *Cx. annulirostris* mosquitoes (7-12 days post eclosion) were starved of sugar and water for 24 hours before being challenged with a meal of chicken blood spiked with JEV supernatant (∼10^7 TCID_50_/ml final concentration) using the Hemotek system. Mosquitoes were left to feed for 1 hour and then anesthetized using carbon dioxide (CO_2_). Blood-engorged females were selected and housed in Plexiglas cages, with 10% sugar ad libitum for a 14-day extrinsic incubation period as previously described [41].

At 4 days post-infection (dpi), a subset (n=30) of mosquitoes were removed from each cage, anesthetized with CO_2_, midguts dissected and transferred to microfuge tubes containing 30 µL PBS (phosphate buffered saline). At 14 dpi, another subset (n=30) of mosquitoes were anesthetized with CO_2_ and legs and wings removed. The mosquito proboscis was inserted into a 20 µL pipette tip filled with 10 µL PBS. Mosquitoes were left to salivate for 20 minutes after which the saliva was washed from the pipette into an additional 10 µL PBS in a microfuge tube. Mosquito carcasses were then individually placed in microfuge tubes with 100 µL PBS. A subset (n=10) of mosquitoes were fixed in 10% neutral buffered formalin (Australian Biostain) at 0, 4 and 14 dpi for immunohistochemistry (IHC) analysis.

### 2.4 Vertical Transmission

In a separate experiment, *Cx. annulirostris* mosquitoes were challenged with a blood meal spiked with JEV NSW/22, as described above. Forty blood-engorged females (P1) were housed in a Plexiglas cage and provided 10% sugar ad libitum. At 3 dpi, a shallow water filled container was provided for mosquitoes to lay egg rafts. Between 4-6 dpi rafts of approximately 200 eggs per raft were frozen at −80°C (E1) or hatched to adulthood (F1). At 6 dpi, P1 mosquitoes were starved of sugar and water and provided a non-infectious blood meal on day 7. Another container was provided at 10 dpi for egg raft collection between 11-13 dpi and rafts were frozen at −80°C (E2) or hatched to adulthood (F2). At 14 dpi, surviving P1 mosquitoes were collected individually in 100 µL PBS molecular analysis. Adult progeny (F1, F2) were collected up to 10 days post-eclosion and pooled (n=10 mosquitoes per pool) for molecular analysis.

### 2.5 Immunohistochemistry

Female mosquitoes exposed to JEV GIV NSW/22 isolate were tested for the presence of viral antigen via IHC using the flavivirus group-reactive Mab 4G4 raised against the Murray Valley encephalitis virus NS1 protein at a dilution of 1:100 [42].

Following the application of the primary antibody, EnVision FLEX+ Mouse Linker (Dako) was applied to amplify the signal. A HRP-labelled secondary antibody, EnVision FLEX/ HRP (Dako), was then applied with PolyDetector Liquid DEC (Bio SB). Prior to IHC staining, sections were antigen retrieved using the Dako PT link and EnVision FLEX Target Retrieval Solution High pH (Dako) for 30 min at 97 °C. A 3% hydrogen peroxide (VWR) solution in EnVision FLEX Peroxidase Blocking reagent (Dako) was also used to eliminate endogenous peroxidase activity within the mosquitoes. Sections were digitised using Pannoramic Scan II (3DHISTECH Ltd) whole slide imager and examined using Xplore software (Phillips). Photomicrographs were taken using the image capture function in Xplore.

### 2.6 Nucleic acid extraction and quantitative reverse transcription PCR (RT-qPCR)

Whole mosquitoes, carcasses and midguts were homogenized in PBS (300 µL total volume) with carbide shards (Daintree Scientific) in a bead beater (MP Biomedicals) at 6.5m/s for 45 seconds for 3 cycles, with samples chilled on ice between cycles. Mosquito pools were homogenised in PBS (500 µL total volume) using the same conditions. Mosquito egg rafts were homogenised individually in 100 µL PBS at 6.5m/s for 45 seconds for 2 cycles, with samples chilled on ice between cycles. Homogenised egg rafts were pooled with ∼600 – 1200 eggs per microfuge tube. All homogenates were centrifuged at 5000 rpm to produce a clarified supernatant. The mosquito saliva sample volume was increased to 75 µL total volume with PBS and was not homogenized.

Fifty microlitres of clarified homogenate or saliva samples were used for nucleic acid extraction using the MagMax 96 Viral RNA Kit (ThermoFisher Scientific) in a MagMAX Express 96 Magnetic Particle Processor (ThermoFisher Scientific), following the manufacturer’s instructions. Ribonucleic acid (RNA) was eluted in 90 µl of elution buffer. RT-qPCR was performed using a JEV-specific assay, based on Shao et al [43], targeting a 63 base pair (bp) nucleotide region of the NS1 gene. Fifteen microlitre reactions contained 5 μL of RNA, 7.5 μL of AgPath One-step RT-PCR buffer (Ambion), 0.6 μL of 25X reverse transcriptase (Ambion), 0.6 μL of 10 μM each primer (forward: 5’-GCCACCCAGGAGGTCCTT; reverse: 5’-CCCCAAAACCGCAGGAAT) (Integrated DNA Technologies), 0.6 μL of 5 μM TaqMan probe (5’-FAM-CAAGAGGTGGACGGCC-MGB) (ThermoFisher Scientific) and 0.1 μL of nuclease free water. The following cycling conditions were used: 10 min at 45 °C for reverse transcription, 10 min at 95 °C for inactivation of RT, followed by 45 cycles of 95 °C for 15 s, 60 °C for 45 s using a 7500 Real-time PCR system (Applied Biosystems) or QuantStudio 5 Real-time PCR system (ThermoFisher Scientific).

A second reference gene RT-qPCR targeting a 106 bp nucleotide region of the *Cx. annulirostris* 18s ribosomal RNA gene was performed on whole mosquitoes, pooled mosquitoes, midgut, carcass and eggs for data normalisation. Fifteen microlitre reactions contained 5 μL of RNA, 7.5 μL of AgPath One-step RT-PCR buffer (Ambion), 0.6 μL of 25X reverse transcriptase (Ambion), 0.3 μL of 45μM each primer (5’-GTGCGTACGGTAGAGAGACAGAGA; reverse: 5’-CGCGCTCGCTCGGTAT) (Integrated DNA

Technologies), 0.3 μL of 12.5 μM TaqMan probe (5’-Cy5-CTAGGCTGG/TAO/TCAGGTCCGGATCGC-IAbRQSp) (Integrated DNA Technologies) and 1 μL of nuclease free water, and as was performed using the same conditions as the JEV-specific assay.

To determine the absolute number of genomic copies of JEV and 18s gene in each mosquito sample, synthetic JEV RNA or *Cx. annulirostris* 18s ribosomal RNA (Integrated DNA Technologies) was designed based on target sequences and used to generate standard curves. Briefly, serial 10-fold dilutions of RNA were prepared in Tris-EDTA buffer and each dilution was tested via RT-qPCR. The limit of detection was 120 JE viral genomic copies and 12 genomic copies for 18s. Linear equations were derived from the standard curves, enabling normalized JE viral genomic copy numbers to be calculated from C_T_ values obtained from RT-qPCR testing of JEV-infected samples. Statistical analysis was performed using GraphPad Prism (Version 9), using unpaired t-test and considered significant when p<0.05.

## 3. Results

### 3.1 Vector competence

To assess the vector competence of *Cx. annulirostris* mosquitoes for the JEV GIV NSW/22 isolate, adult female mosquitoes were exposed to chicken blood spiked with JEV (∼10^7 TCID_50_/ml). Results from RT-qPCR testing of mosquito midguts (n=29) dissected at 4 dpi showed a 100% infection. At 14 dpi, RT-qPCR of mosquito carcasses (n=30) showed a 100% dissemination and an 87% positivity rate for the virus in saliva ((Table 2; Figure 1). Together, these results demonstrate that *Cx. annulirostris* is an efficient vector of the NSW/22 isolate.

**Table 2:** Infection, dissemination, and transmission rates in *Cx. annulirostris* mosquitoes exposed to JEV isolates.

| JEV isolate | % Infection<sup>a</sup> | % Dissemination<sup>b</sup> | % Transmission<sup>c</sup> |
| --- | --- | --- | --- |
| NSW/22 | 100 (29/29) | 100 (30/30) | 87 (26/30) |
| TS00 | 87 (26/30) | 100 (30/30) | 53 (16/30) |
| FU | 73 (22/30) | 97 (29/30) | 83 (25/30) |
| JKT6468 | 76 (22/29) | 100 (30/30) | 70 (21/30) |

<sup>a</sup> Percentage of mosquitoes with JEV in the midgut (number positive/number tested)
<sup>b</sup> Percentage of mosquitoes with JEV in the carcass (number positive/number tested)
<sup>c</sup> Percentage of mosquitoes with JEV in the saliva (number positive/number tested)

**Figure 1.**
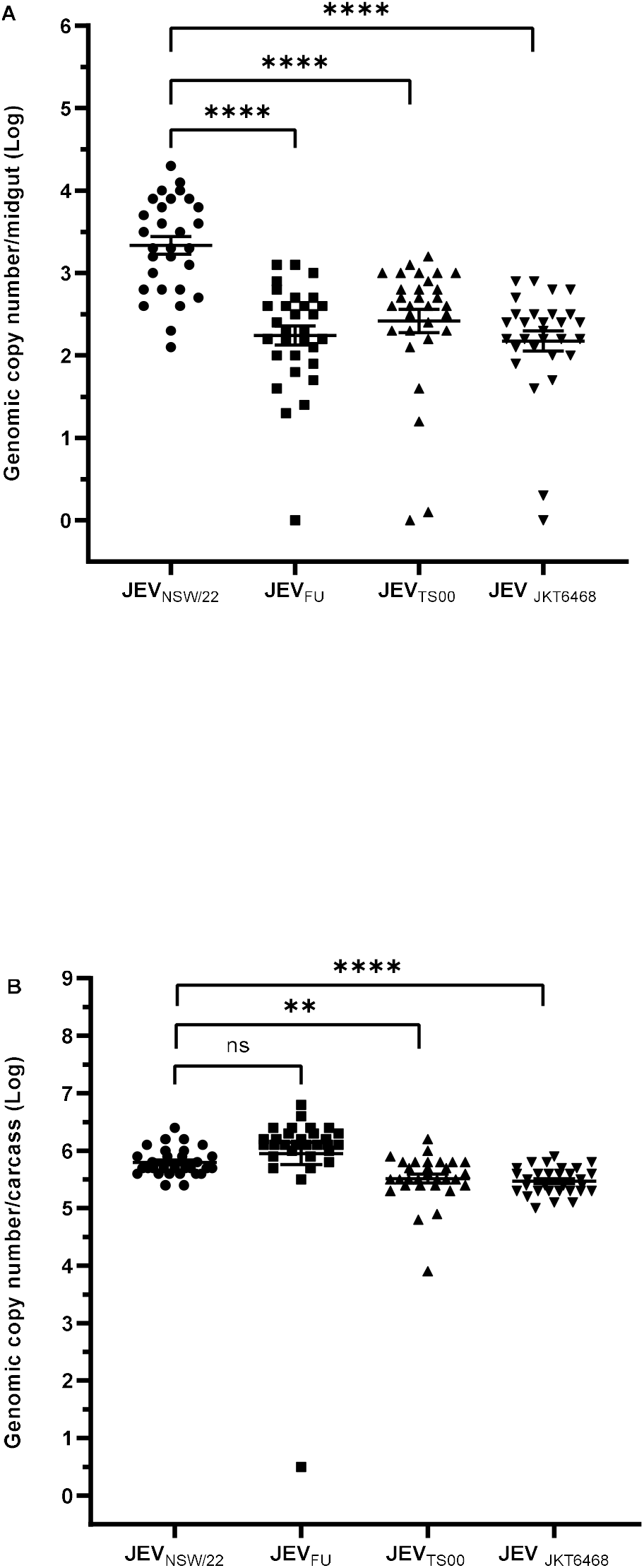
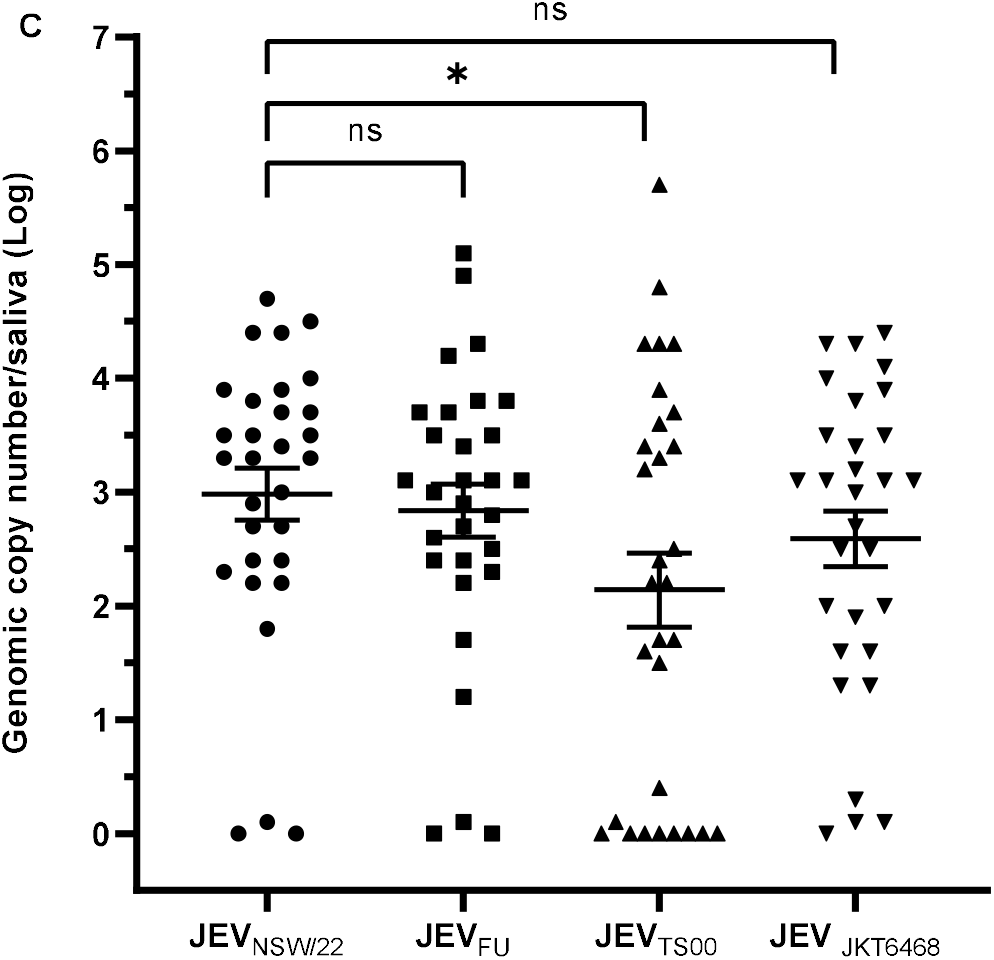
Vector competence of *Cx. annulirostris* mosquitoes for JEV isolates. *Cx. annulirostris* mosquitoes were artificially infected with JEV isolates as described in Methods. JEV-specific real-time RT-qPCR was performed on midguts (A), carcass (B), and saliva (C). *Cx. annulirostris* 18S real-time RT-qPCR was performed for normalisation and a standard curve using synthetic RNA was used to calculate genomic copy numbers. Each dot denotes individual mosquitoes and horizontal bars denote the mean with standard error of the mean. ns, not significant; significant differences: *, p<0.05; **, p<0.005; ****, p<0.0005.

For comparison, adult female *Cx. annulirostris* mosquitoes were exposed to previous Australian isolates of JEV belonging to genotypes I (TS00) and II (FU), as well as an early GIV isolate from Indonesia (JKT6468). Mosquitoes were artificially fed blood spiked with the individual virus (∼10^7 TCID_50_/ml) and results of RT-qPCR analysis showed high midgut infection (≥73%) and high carcass dissemination (≥97%) rates for these viruses (Table 2). High levels of virus positivity (83%) in saliva were found in JEV FU infected mosquitoes, while mosquitoes infected with JEV TS00 and JKT6468 showed 53% and 70% virus positivity in saliva respectively. These results suggest that *Cx. annulirostris* is an efficient vector for all tested genotypes of JEV, with variation in infection, dissemination and transmission rates observed between tested isolates. Interestingly, at 4 dpi to JEV NSW/22, there were statistically significantly higher mean viral genome copies in the mosquito midguts compared to those of mosquitoes exposed to TS00, FU and JKT6468 isolates (p=<0.0005). At 14 dpi, there were also statistically significantly higher mean viral genome copies detected in the carcasses of JEV NSW/22 exposed mosquitoes compared to TS00 (p=<0.005) and JKT6468 (p<0.0005). However, there were no significant differences in viral copies in the carcass and saliva of JEV NSW/22 compared to FU (p>0.05) (Figure 1).

### 3.2 Immunohistochemistry

To determine the presence and tropism of JEV NSW/22 in infected mosquitoes, IHC was performed on whole mosquito sections at 0, 4 and 14 dpi (Figure 2a-o). Viral NS1 antigen was detected at 4 dpi, predominantly in the mid-gut (Figure 2e), and to a much lesser extent in rare haemocyte-like cells in the stroma of the ovary (Figure 2k) and adjacent to the head ganglion (Figure 2n). At 14 dpi, NS1 antigen was detected throughout the body, including the midgut epithelium (Figure 2f), salivary gland epithelium (Figure 2i), neurons of the head ganglion (Figure 2o, arrowhead), photoreceptor cells of the compound eye (Figure 2o, arrow), cells associated with the basilar aspect of the hindgut and Malpighian tubules, and haemocyte-like cells in ovarian stroma (Figure 2l). Notably, NS1 antigen was not detected in germ cells of the ovary.

**Figure 2.**
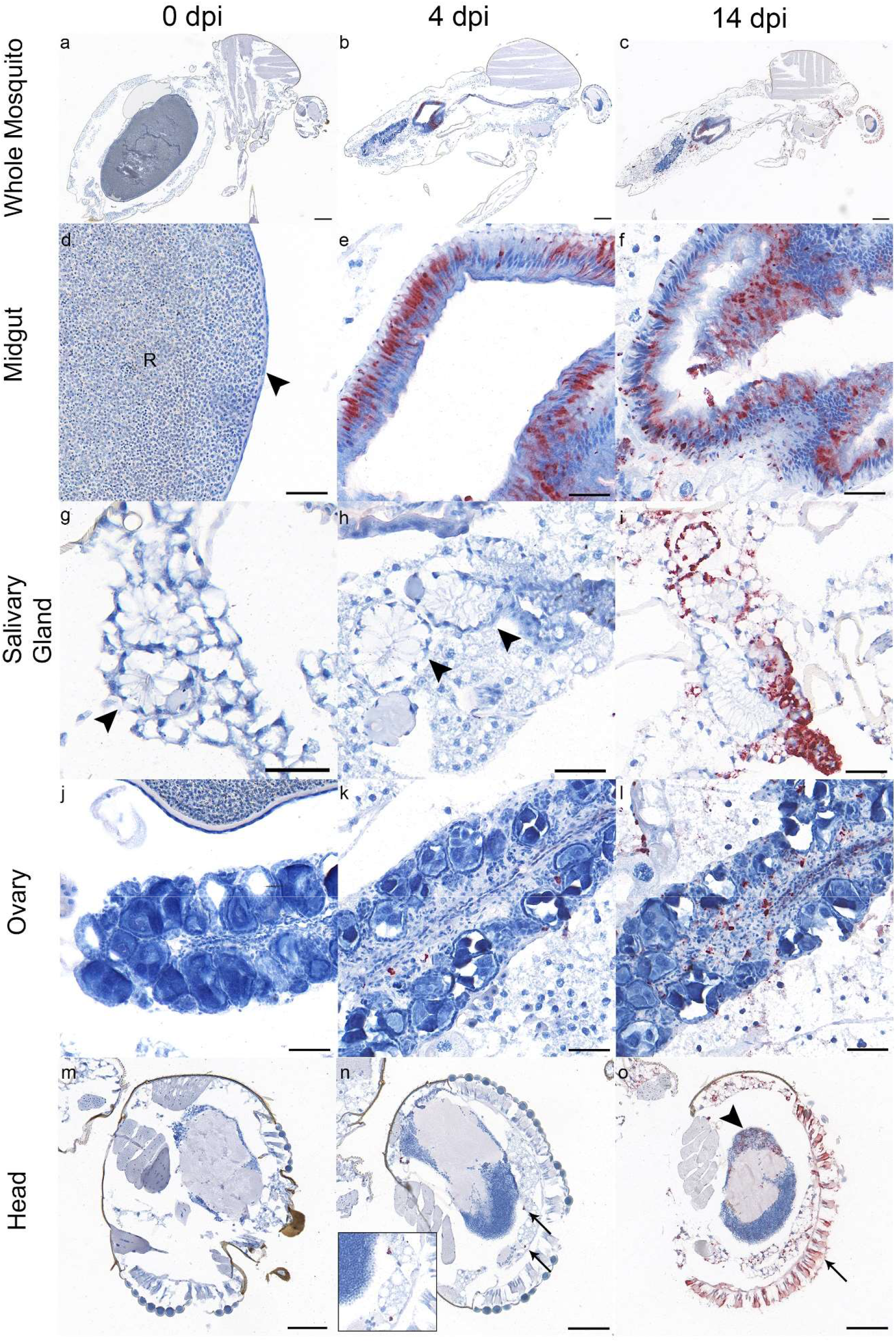
Tropism of GIV JEV isolate NSW/22 in *Cx. annulirostris* mosquitoes infected via artificial bloodmeal on 0 (left column), 4 (middle column), and 14 dpi (right column), in the midgut (panel d-f), salivary gland (panel g-i), ovary (panel j-l), and head (panel m-o). Red labelling represents the presence of viral NS1 antigen. Antigen was detected in midgut epithelium from 4 dpi (panel e and f); salivary gland epithelium on 14 dpi (panel i); haemocyte-like cells in ovarian stroma from 4 dpi (panel k and l); haemocyte-like cells adjacent to the head ganglion on 4 dpi (panel n, arrows; inset is an enlarged image of the antigen positive cells indicated by the arrows), and head ganglion neurons (panel o, arrowhead) and photoreceptor cells of the compound eye (panel o, arrow) on 14 dpi. In panel d, “R” denotes ingested red blood cells in the lumen of the midgut, and arrowhead indicates the severely flattened midgut epithelium after blood meal. In panels g and h, the arrowheads indicate acini of the salivary gland. Scale bars: panel a-c: 250 µm; panel d-f: 50 µm; panel g-l: 50 µm; and panel m-o: 100 µm.

### 3.3 Vertical Transmission

Experiments were conducted to determine whether *Cx. annulirostris* mosquitoes can transmit JEV NSW/22 to their progeny. RT-qPCR testing completed on extracts of egg rafts laid after first (E1, JEV NSW/22) and second (E2, non-infective) blood meals detected the presence of JE viral RNA in 100% of E2 raft pools, indicating transovarial transmission of virus. However, no JE viral RNA was detected in adults reared from these egg rafts indicating that trans-stadial transmission of JEV NSW/22 did not occur from infected *Cx. annulirostris* (Table 3).

**Table 3:** Vertical transmission of JEV NSW/22 in *Cx. annulirostris* mosquitoes.

| Life stage | % Positive |
| --- | --- |
| E1 egg rafts | 33 (1/3)<sup>a</sup> |
| F1 mosquito progeny | 0 (0/35)<sup>b</sup> |
| E2 egg rafts | 100 (3/3)<sup>c</sup> |
| F2 mosquito progeny | 0 (0/13)<sup>d</sup> |

<sup>a</sup> Percentage of positive egg raft pools (E1) after first infectious bloodmeal (number positive/number tested)
<sup>b</sup> Percentage of positive mosquito progeny pools (F1) reared from E1 (number positive/number tested)
<sup>c</sup> Percentage of positive egg raft pools (E2) after second non-infectious blood meal (number positive/number tested)
<sup>d</sup> Percentage of positive mosquito progeny pools (F2) reared from E2 (number positive/number tested)

## 4. Discussion

During the 2022 Australian outbreak of JEV genotype IV, the virus was detected in field-collected *Cx. annulirostris* [37]. While previous laboratory experiments with this mosquito species have been conducted using the most commonly circulating genotypes (I-III) [32], the role of *Cx. annulirostris* in the transmission of JEV GIV using experimental infections remained to be confirmed. Our results showed that Australian *Cx. annulirostris* is a highly competent vector of the 2022 Australian outbreak strain of GIV JEV, with 87% of mosquito saliva samples from infected mosquitoes testing positive for JEV, indicative of high transmission potential (Table 2; Figure 1). This virus was also shown to be capable of widespread dissemination within the tissues of infected *Cx. annulirostris*, as demonstrated by IHC analysis.

For comparison with experiments performed using the JEV NSW/22 isolate, vector competence experiments were also performed with representative isolates of JEV, including a progenitor GIV isolate from Indonesia and isolates of genotypes previously detected in Australia[12,19,39], JEV genotype II emerged in the Torres Straits of Australia in 1995[12] and in 2000, GI JEV (later shown to be G1b) [9] detected for the first time in the Torres Strait [39], apparently displacing GII viruses, with no further GII detections subsequently made in northern Australia. Vector competence experiments with these isolates in our study showed that, although levels of JEV detected in the saliva of JEV JKT6468 and FU infected mosquitoes were comparable to NSW/22 infected mosquitoes, only 53% of saliva samples from mosquitoes infected with JEV TS00 strain were positive and contained significantly lower levels of virus (Table 2, Figure 1). This suggests that the transmission potential of *Cx. annulirostris* infected with JEV TS00 may be lower than that following infection with other isolates. While this finding is inconsistent with the displacement of GII viruses by JEV GI in northern Australia, our data may explain the displacement of GI with the novel lineage of JEV GIV in northern Australia. A previous vector competence study using a lab colony of *Cx. annulirostris* at 14 days post-infection with a 1998 GII Torres Strait isolate (TS3306), showed almost identical transmission rates (81%); however, in contrast to our findings, dissemination was much lower (64%) [32]. Another study also found levels of infection with the GI TS00 at 12 dpi comparable with our study (80%); however, dissemination (56%) and transmission (12%) were considerably lower [30]. The differences between the results of these studies and those presented here may be due to the different colonies of *Cx. annulirostris* employed and virus detection methods used.

Notably, significantly higher viral titres were observed in the midguts of JEV NSW/22 infected mosquitoes at day 4 post-infection in comparison to the other isolates tested (Table 2; Figure 1). By day 14, there was no significant difference in the viral genomic copy numbers of JEV NSW/22 and FU detected in mosquito carcasses; however viral copy numbers remained significantly higher for mosquitoes infected with JEV NSW/22 compared to TS00 and JKT6468. These genotype and/or strain-specific differences are not surprising, as they have previously been reported for *Cx. quinquefasciatus* [44]. This study showed significantly higher infection rates in mosquitoes exposed to GI-b and GIII strains compared to a GI-a strain, suggesting mosquito vector competence could play a role in the selective advantage of one genotype over another and lead to the maintenance of that genotype in the enzootic transmission cycle.

Vertical transmission of JEV from infected females to their progeny after overwintering has been shown in various mosquito species with varied efficacy, including *Cx. tritaeniorhynchus* [34]. Our results showed efficient transovarian transmission of JEV NSW/22; however, there was no trans-stadial transmission from eggs to adults observed (Table 3). The IHC results provided a potential explanation for this observation: JEV replication was detected in haemocyte-like cells in the ovarian stroma rather than in germ cells (Figure 2). This may be responsible for the egg rafts testing positive but without virus transfer to the mosquito progenies reared from the eggs. Further experiments, including transmission studies at different temperatures and with field-caught mosquitoes, need to be performed to determine the role of overwintering in the persistence of JEV GIV in *Culex* mosquitoes.

Our study provides evidence that Australian *Cx. annulirostris* mosquitoes are highly competent for the Australian outbreak strain of JEV GIV. The relatively higher levels of virus detected in the midgut and saliva indicate that this strain replicates more efficiently in and is transmitted by the major vector species of JEV in Australia and provides a plausible mechanism by which this strain emerged in Australia as the dominant genotype of JEV. Further studies are needed to determine the vector competence of other mosquito species found in Australia that have been implicated as vectors of JEV, such as *Cx. quinquefasciatus, Cx. sitiens, Cx. gelidus* and *Cx. tritaeniorhynchus*, to inform the risk of sustained transmission of JEV in Australia. Vector competence of different populations of the same species can also vary [32], therefore it will be important to test a range of populations of the same mosquito species, especially collected from the field, to determine regional/geographic differences and to inform the risk of spread into other regions. There is also the need for continued surveillance in countries neighbouring northern Australia, including Indonesia, PNG, and Timor Leste, to understand the level of diversity of circulating strains of JEV and their potential to emerge and displace existing genotypes. Whether this JEV GIV will displace currently circulating genotypes in the Indo-Pacific countries also remains to be seen.

Various environmental and genetic factors, such as weather and host susceptibility, may play a role in JEV emergence. Our data suggests that increased infectivity for the outbreak strain, indicated by comparatively higher levels of virus replication and transmission rate from saliva may have contributed to the apparent rapid spread of the JEV across Australia.

## Declarations

### Author contribution statement

MJK contributed to the design of the study, data acquisition, analysis, interpretation and drafting of the manuscript. SJ and KRB were involved in the planning and data acquisition. MD and AJLD were involved in data acquisition. DB, MH, DG and JW performed the molecular analysis of the samples. WWS and JP performed the immunohistology and interpretation of the images. DTW and PNP contributed to the overall study concept, design, data interpretation and manuscript drafting. All authors contributed to the writing and review of the manuscript.

## Acknowledgements

We thank Dr. Peter Kirkland (Elizabeth Macarthur Agriculture Institute, NSW) for providing the JEV GIV-O-0883/NSW/2022 isolate. We acknowledge the use of the CSIRO Australian Centre for Disease Preparedness (https://ror.org/02aseym49) in undertaking this research. We would also like to thank Dr. Helle Bielefeldt-Ohmann (University of Queensland) for her insights into mosquito histoanatomy and IHC of flaviviruses in mosquitoes.

## Ethics Approval

ACDP Animal Ethics 1957, 22010

## Data Availability statement

All relevant data is available in the article.

